# Mechanical polarity links adhesion-tuned protrusions to directional stability in glioblastoma cell migration

**DOI:** 10.1101/2025.10.23.684087

**Authors:** Haruna Tagawa, Daisuke Kanematsu, Asako Katsuma, Naoyuki Inagaki, Yonehiro Kanemura, Yuichi Sakumura

## Abstract

Glioblastoma invasion critically limits therapeutic outcomes, and understanding the physical principles that govern cell motility is essential for developing effective therapies. Here, to clarify the mechanical links between cell adhesion, protrusions, and migration, we analyzed glioblastoma-derived cells migrating on fibronectin- and laminin-coated extracellular matrix (ECM) substrates using time-lapse imaging and mathematical modeling. We quantified cell motility on each ECM and constructed a coarse-grained biophysical model that incorporates catch- and slip-bond kinetics. We treated the ECM as an external boundary condition that modulates adhesion dynamics, distinct from the intrinsic mechanical parameters of the cell. The model reproduced ECM-dependent differences in cell motility. The results suggest that adhesions and protrusion elongation on fibronectin are less stable than on laminin, and opposing forces between protrusions reduce net displacement more strongly on fibronectin than on laminin, leading to less coordinated tensile forces. Based on these results, we establish *mechanical polarity*, defined as an imbalance of protrusion- and adhesion-mediated forces that drive directional migration, as a quantifiable physical principle of cell protrusion and migration. This principle likely extends beyond glioblastoma biology, providing a generalizable mechanism that links adhesion dynamics to migration stability and offering a physical basis for strategies to suppress invasive cell behavior.

## Introduction

Tumor cell invasion is a central driver of cancer progression, and deciphering the mechanisms of uncontrolled cell migration remains a critical challenge in tumor biology. Gliomas are the most common adult primary brain tumors, and more than half of them are glioblastomas, which continue to be associated with dismal patient outcomes (1–4). The highly invasive motility of glioblastoma cells underlies this therapeutic intractability.

Glioblastoma cells extend protrusions called tumor microtubes (5, 6) that dynamically attach to and detach from the extracellular matrix (ECM), potentially governing both migration efficiency and directional stability. However, the physical mechanisms, including how cellular forces and adhesion dynamics interact with the ECM and reflect intrinsic motile properties, have not been fully elucidated. To approach this problem from a physical perspective, we consider the ECM as an external boundary condition that modulates adhesion dynamics, with potential biochemical signaling implicitly embedded in these interactions.

Cell migration is an integrated phenomenon that arises from morphological polarization, cytoskeletal remodeling, and adhesion dynamics. Traditionally, directional stability of migration has been explained by morphological asymmetry (7–10). However, some observations suggest that cells can maintain stable migration directions even when morphological polarity is weak or misaligned with the direction of movement (11, 12), challenging the conventional framework that links shape asymmetry to directional persistence.

These observations highlight a missing link between cellular morphology and the mechanics of persistent migration. Understanding such behavior requires identifying the physical basis of directional stability. Yet how the imbalance between substrate traction and the corresponding reaction forces transmitted through protrusions, referred to here as protrusive forces, governs persistent migration has not been quantitatively validated in living cells. Here, we define *mechanical polarity* as the physical imbalance of these protrusive and adhesive forces that underlies directional stability in glioblastoma cell migration.

While most existing mathematical models describe cells as rigid or continuous bodies without explicitly representing local protrusive dynamics (13, 14), the Cell Migration Simulator (CMS) (15) successfully incorporated protrusion formation and force transmission to establish a detailed mechanical framework for cell migration. Building on this foundation, we developed a complementary, coarse-grained model that integrates protrusion dynamics, adhesion kinetics, and molecular transport to quantify how force balance governs cell migration.

To overcome the computational and conceptual limitations of detailed models, we employed the glioblastoma-derived cell line GDC40 (16) under serum-free conditions that better reflect *in vivo*–like intrinsic motility, and performed time-lapse analyses of cell migration on fibronectin- and laminin-coated substrates to quantify protrusion and cell body dynamics. This design allowed us to isolate cell-intrinsic mechanical parameters by comparing motility under distinct ECM environments. Based on these data, we developed a coarse-grained mathematical model that incorporates the molecular characteristics of catch and slip bonds. Whereas a slip bond is an interaction in which force accelerates bond dissociation and thereby shortens bond lifetime, a catch bond is an interaction in which force prolongs bond lifetime (17–19). Our model enables efficient simulation of protrusion-dependent adhesion–detachment dynamics, recapitulates ECM-dependent differences in motility, and reveals that in cells on fibronectin, weaker adhesion stability leads to uncoordinated protrusive forces, causing opposing forces between protrusions to reduce migration efficiency more than in cells on laminin.

Although it is physically evident that a cell moves in the direction of greater net force, this relationship has rarely been quantified in living cells where protrusion–adhesion interactions are highly dynamic. These findings demonstrate that in glioblastoma cells, directional stability arises from the imbalance of protrusive and adhesive forces, referred to here as mechanical polarity, rather than from morphological asymmetry.

## Results

### Glioblastoma cell migration depends on ECM coating conditions

We analyzed time-lapse images of GDC40 cells on glass coated with fibronectin (F-GDC40) (**Fig. 1A, Movie S1**) or laminin (L-GDC40) (**Fig. 1B, Movie S2**). From the images, we extracted the coordinates of the cell body center and protrusion tips (**Fig. 1C**), and quantified two groups of features: migration (turning angle and speed; **Fig. 1D**) and protrusion features (number, maximum length, and lifetime; **Fig. 1E**).

F-GDC40 frequently changed direction by more than 90° (**Fig. 1F**). In contrast, L-GDC40 mostly turned by less than 90°, indicating higher directional persistence. L-GDC40 showed greater migration speed, longer protrusions, and fewer protrusions than F-GDC40. No significant difference was found in protrusion lifetime. Since cell migration depends on protrusion dynamics (10, 20), these results suggest that the differences in protrusion features that we observed can substantially affect migration features.

### Development of a quantitative model of the cell migration

Protrusion dynamics are mechanically regulated by adhesion and detachment events at the protrusion tip, as well as changes in tensile forces (21). We hypothesized that the differences in the mechanical properties of adhesion could alter protrusion number and length, thereby shaping the migration phenotype. To test this, we developed a quantitative physical model based on our previously reported mathematical model of neuronal polarization (22) and conducted simulations.

The model represents a glioblastoma cell as a central cell body with multiple protrusions (**Fig. 2A**). The cell body migrates according to the vector sum of forces generated by the protrusions (**i**). Protrusions emerge stochastically (**ii**) and are retracted and removed when their length falls below a defined threshold (**iii**). Clutch molecules reversibly link adhesion molecules to the actin cytoskeleton; when engaged, they couple retrograde actin flow to the ECM to generate traction and, by reaction, protrusive force, whereas disengagement releases these forces (23–25). We therefore hypothesized that protrusion elongation depends on the transport of clutch molecules from the cell body to the tip and their return by diffusion as observed in neuronal cells (26) (**iv**), as well as on the balance between protrusive force and tension at the tip (**v**). The effectiveness of the adhesion is represented by the bound fraction *g*, defined as the ratio of the number of adhesion molecules bound to the ECM to the total adhesion molecules at the protrusion tip, modeled as a function of protrusive force (**vi**). Further details of the model are provided in the Materials and Methods.

**Fig. 1.**
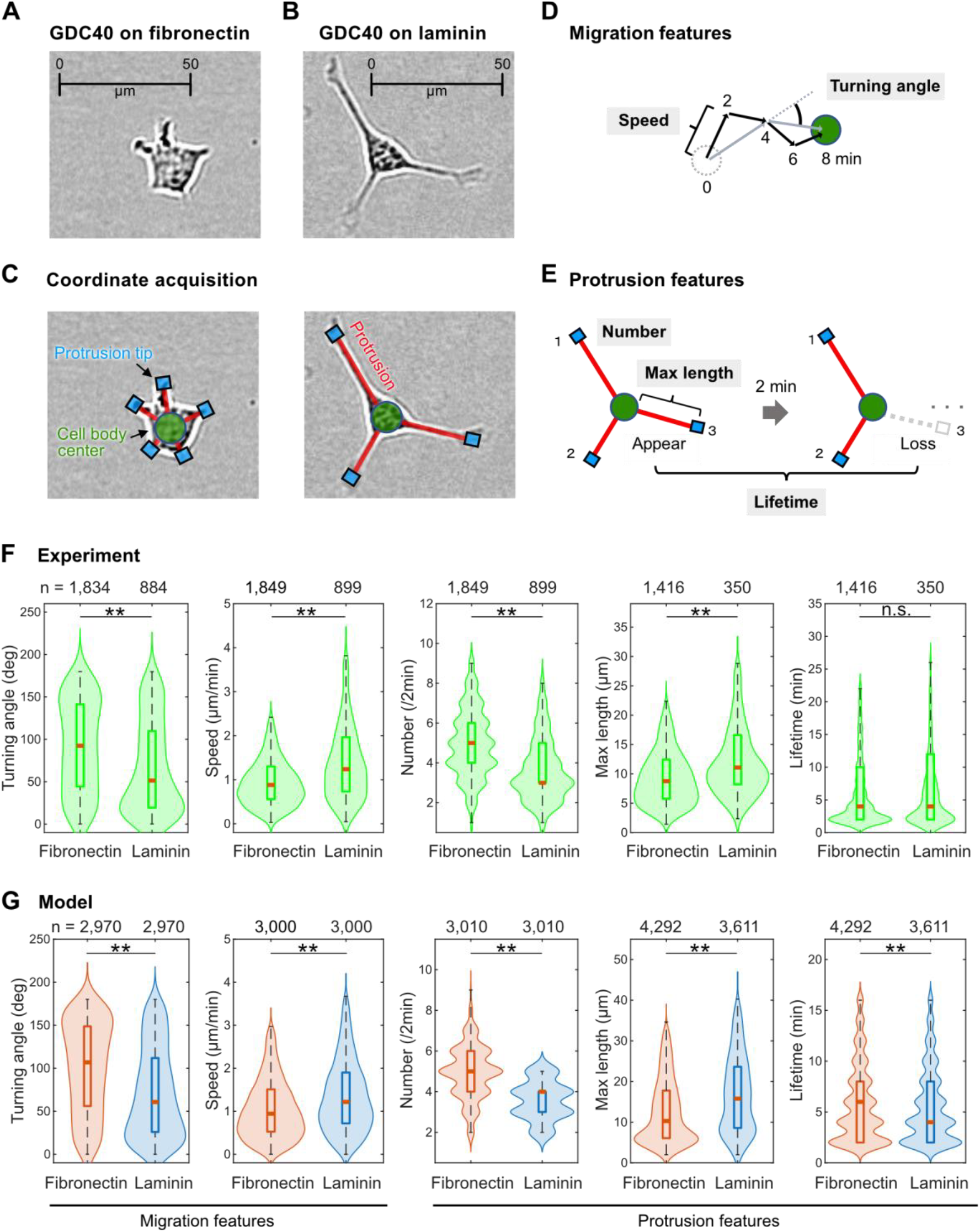
Quantitative features of GDC40 cell migration and protrusion dynamics. (A, B) Phase-contrast image of a GDC40 cell on fibronectin and on laminin. (C) Quantified components: cell body center (green), protrusion tips (cyan squares), and protrusion lengths (red lines). (D) Definitions of migration features. Speed was defined as the displacement of the cell body center over 2-minute intervals, whereas the turning angle was defined as the change in migration angle over 4-minute intervals to reduce the effect of short-term fluctuations. (E) Protrusion features include the number per frame, maximum length, and lifetime. (F, G) Quantification results of migration and protrusion features for GDC40 cells on fibronectin (F-GDC40) or laminin (L-GDC40), using five cells per ECM condition. (F) shows results obtained from actual cell images, while (G) shows results from mathematical model simulations. The sample size *n* is indicated above each plot. The red line in the box represents the median. Values more than 1.5 times the interquartile range away from the top and bottom of the box were considered outliers. The outliers were omitted from the plots for visibility, but all data were included in the statistical tests and analyses. Statistical differences were assessed using the Wilcoxon rank-sum test (**p < 0.01; n.s., not significant).

**Fig. 2.**
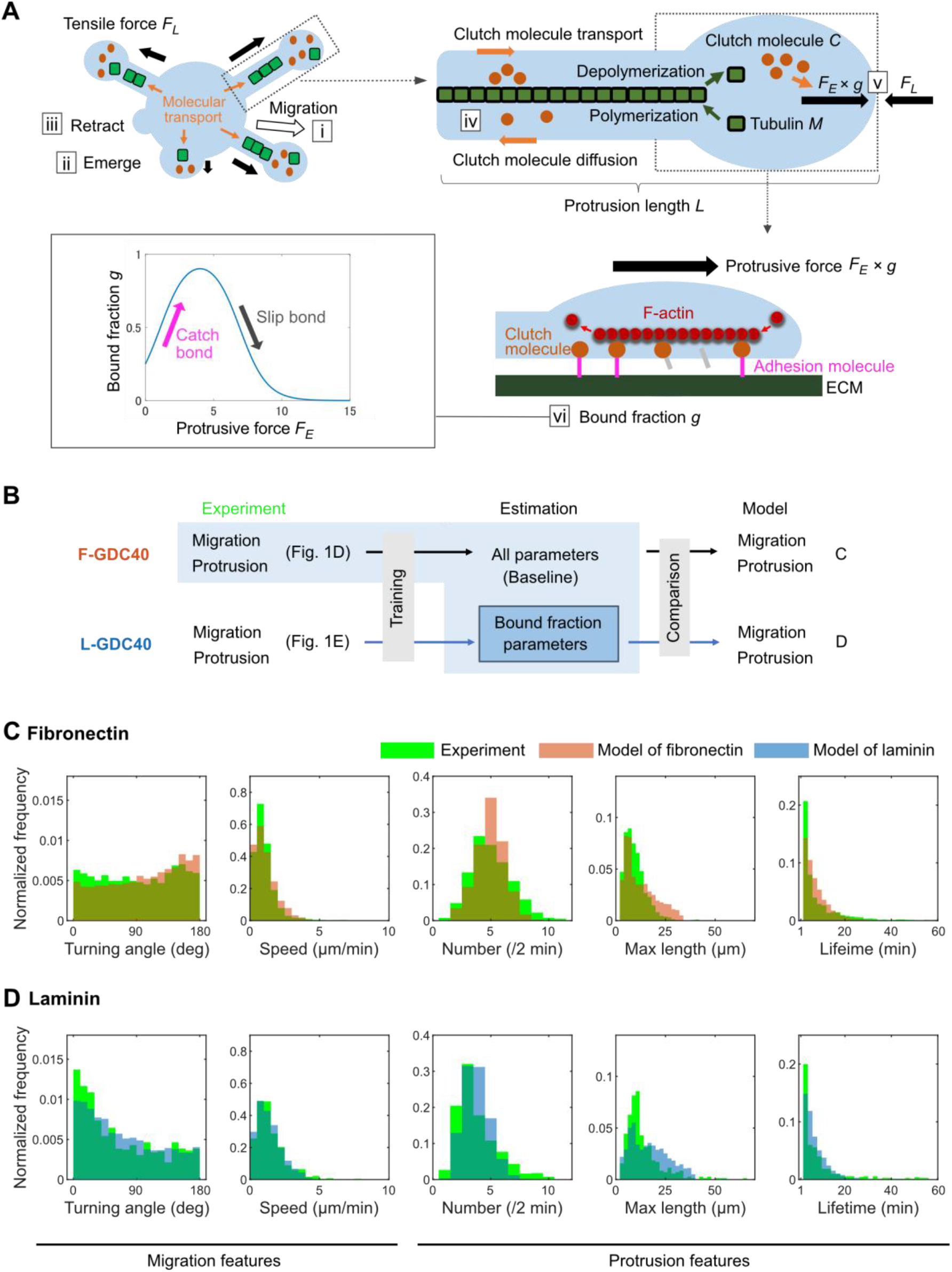
Model construction, parameter fitting, and reproduction of GDC40 cell motility. (A) Schematic of the cell migration model: (i) whole-cell migration driven by protrusion forces; (ii-iii) stochastic emergence and retraction of protrusions; (iv) transport of cytoskeletal components (tubulin) and clutch molecules; (v) force balance at the protrusion tip between polymerization-induced protrusive force *F*_*E*_ *g* and tension *F*_*L*_; (vi) force-dependent bound fraction described by the function *g*(*F*_*E*_), capturing both catch and slip bonds of adhesion molecules. (B) Parameter fitting scheme: All model parameters were first fitted to F-GDC40 data (baseline parameters), followed by selective fitting of the function *g*(*F*_*E*_) to L-GDC40 data. (C, D) Comparison between experimental (green) and simulated (orange or blue) distributions of cell behavior (C: F-GDC40; D: L-GDC40). Sample sizes are shown in **Fig. 1F and G**.

### ECM-dependent migration differences reproduced by adhesion mechanics

To validate the model structure, we conducted parameter estimation and simulations (**Fig. 2B**). We first obtained the feature distributions of F-GDC40 (green distributions of **Fig. 2C**), then estimated all parameters from the distributions and simulated cell migration. We designated this parameter set as the “baseline” (F-ECM condition). Next, using the L-GDC40 feature distributions (green distributions of **Fig. 2D**), we re-estimated only the four adhesion-related parameters (*a*_*s*_, *b*_*s*_, *a*_*c*_, and *b*_*c*_), keeping the rest fixed (L-ECM condition) (**Fig. 2B**). The details of the estimated adhesion-related parameters will be discussed in the next section.

The model reproduced ECM-dependent differences in migration and protrusion, including a lower turning angle and higher speed in L-GDC40, as well as more numerous and shorter protrusions in F-GDC40 (**Fig. 1G, Fig. 2C and D, *SI Appendix*, Fig. S1, and Movie S3 and S4**). We calculated the Jensen–Shannon (JS) divergence to compare model feature distributions with experimental data under each ECM condition. A smaller JS divergence indicates greater similarity between distributions. In both ECM conditions, except for lifetime (which showed no significant differences in the experiments; **Fig. 1F**), the JS divergence was smaller for model–experiment pairs with the same ECM type than for non-matching pairs (***SI Appendix*, Table S1**), confirming that each model best reproduced the migration and protrusion features of its respective ECM condition. Furthermore, even when adhesion parameters were estimated using only migration features or only protrusion features, the differences in motility between L-GDC40 and F-GDC40 were still successfully reproduced (***SI Appendix*, Fig. S2 and Table S2**).

Notably, these migration features were reproduced solely by modifying adhesion-related parameters. These results suggest that the differences in migratory behavior between F-GDC40 and L-GDC40, as observed in our experiments, can be explained by mechanical differences in adhesion. Moreover, the fact that these distinct migratory patterns emerged by varying only adhesion-related parameters strongly supports the validity of our model in capturing intrinsic migratory mechanisms of glioblastoma cells.

### Differences in adhesion properties alter mechanical interference between protrusions and modulate cell motility

Our model assumes a causal relationship in which the bound fraction of adhesion molecules regulates protrusion features, which in turn influence migration features. To test this assumption, we analyzed the simulation results to quantify how differences in bound fraction and in the resulting tensile forces between the F-ECM and L-ECM conditions contribute to cell migration.

The estimated ratios of the four adhesion-related parameters (*a*_*s*_, *b*_*s*_, *a*_*c*_, *b*_*c*_) between the F-ECM and L-ECM conditions were 1.07, 1.38, 0.91, and 1.17, respectively. *a*_*s*_, *b*_*s*_, *a*_*c*_, and *b*_*c*_ are compiled parameters of the number of adhesion molecules and adhesion sites on the extracellular matrix, bond strength, reaction rate constants, and the length per unit of the cytoskeletal components (***SI Appendix*, SI Text)**. *a*_*s*_ and *b*_*s*_ are attributable to slip bonds, while *a*_*c*_ and *b*_*c*_ are attributable to catch bonds. *a*_*s*_ and *a*_*c*_ act as sensitivity parameters, and *b*_*s*_ and *b*_*c*_ serve as scale parameters (see Methods, Eq. (7)).

In the F-ECM condition, the lower value of the catch bond sensitivity parameter *a*_*c*_ and the higher values of the scale parameters *b*_*s*_ and *b*_*c*_ indicate that the bound fraction responds more gradually to force and reaches a lower maximum level compared to the L-ECM condition. Consistently, the bound fraction function *g*(*F*_*E*_), computed from these parameters, showed higher values across all force levels *F*_*E*_ in the L-ECM condition than in the F-ECM condition (**Fig. 3A**). Notably, the peak value of *g*(*F*_*E*_) was approximately 0.45 for F-ECM and 0.5 for L-ECM, suggesting that cell–ECM adhesion is more unstable and prone to slippage under the F-ECM condition. Because the bound fraction function exhibits a switch-like shape, the distribution of bound fraction values during model simulations was bimodal under both ECM conditions; the L-ECM condition showed a slightly broader distribution (**Fig. 3B**).

**Fig. 3.**
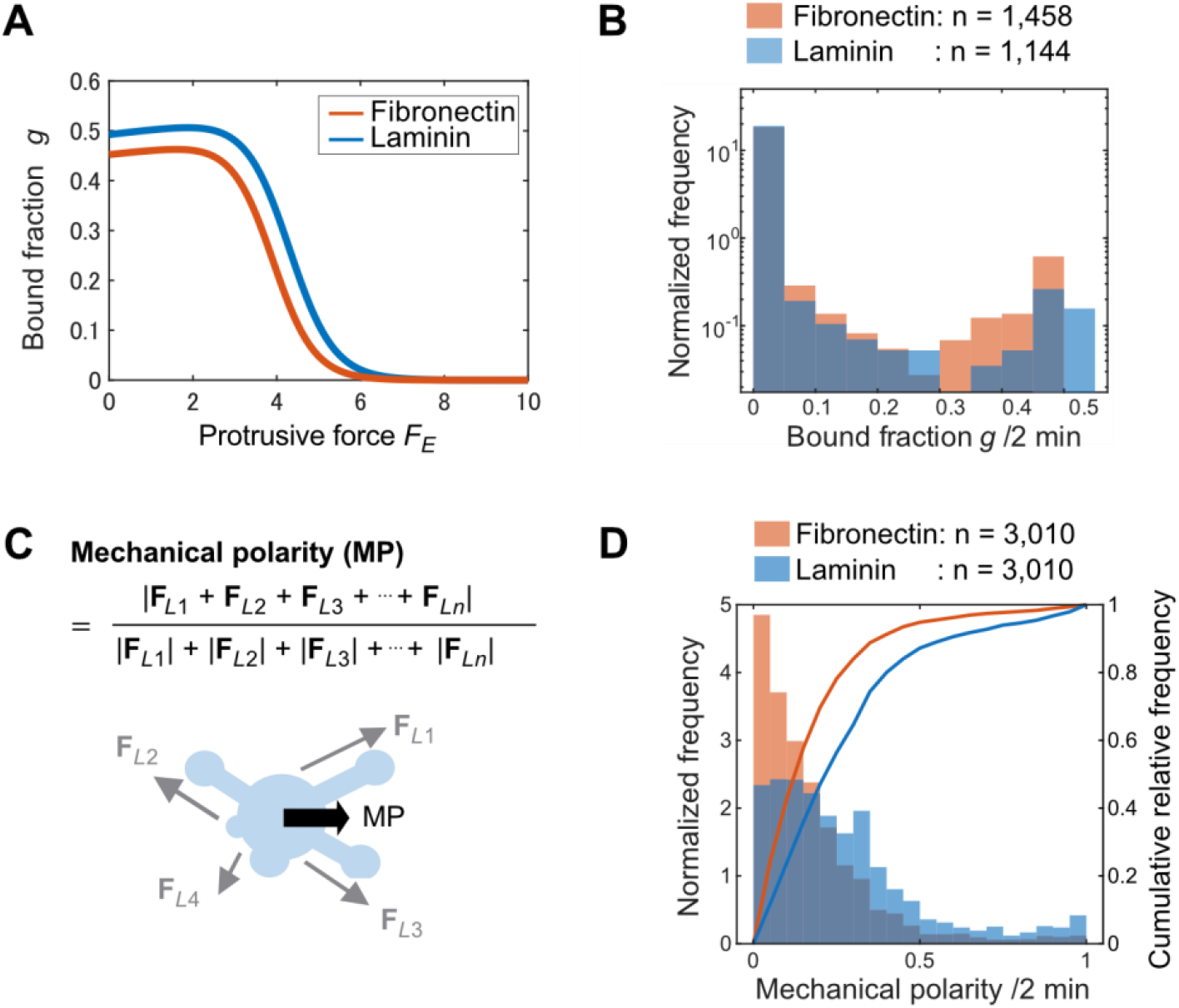
ECM-dependent adhesion dynamics and protrusion force efficiency in the model. (A) The bound fractions *g*(*F*_*E*_) under two ECM conditions. The orange curve represents the baseline parameters estimated from F-GDC40 (F-ECM condition), and the blue curve shows the result of refitting the bound fraction parameters (*a*_*c*_, *b*_*c*_, *a*_*s*_, *b*_*s*_) using L-GDC40 data (L-ECM condition). *F*_*E*_ denotes the protrusive force excluding the effect of the bound fraction. (B) Log-scaled histograms of bound fraction *g* measured every 2 minutes for each protrusion in a single simulated cell. Orange and blue represent F-ECM and L-ECM conditions, respectively. (C) Definition of mechanical polarity as the ratio between the net traction force vector and the total magnitude of all protrusion forces. (D) Distributions and cumulative plots of mechanical polarity obtained from simulations. Values were sampled every 2 minutes over 10 simulated cells per condition. Data points where protrusions were absent or forces were zero were excluded.

To understand how protrusion-generated forces contribute to cell migration, we introduce the concept of mechanical polarity, which represents the forces acting on the cell body that do not completely cancel out but remain biased in a particular direction, thereby driving the displacement of the cell body. Mechanical polarity is defined as the magnitude of the resultant force divided by the sum of the magnitudes of the individual forces (**Fig. 3C**). It quantifies the efficiency of such force generation. A higher value indicates better alignment of forces and more efficient translation of protrusion force into net displacement of the cell body. Polarity, in its original sense, refers to directionality. In this study, polarity is distinguished from biological polarity based on morphological or molecular distributions; instead, it is defined as the directional property arising from non-zero physical forces acting on the cell body.

Simulation results for mechanical polarity are shown in **Fig. 3D**. Values below 0.2 occurred more frequently under F-ECM conditions, while values above 0.2 were more common under L-ECM. The statistical test (Wilcoxon rank-sum test) revealed a significant difference in the values between F-ECM and L-ECM (p < 0.01). This indicates that in L-GDC40, which exhibit higher speed and directional persistence, protrusion forces are less likely to cancel out and more efficiently drive directional migration.

### Protrusion frequency regulates migration efficiency and offers a potential strategy for motility suppression

Based on our prior results, we hypothesized that modulating protrusion features, while keeping adhesion parameters constant, could control mechanical polarity and thereby influence migration features. In particular, converting the “few long protrusions” characteristic of L-GDC40 into the “many short protrusions” observed in F-GDC40 may reduce motility.

To test this hypothesis, we increased the protrusion rate parameter *k*_*pr*_ from its estimated value of 0.134 to 0.2, while keeping all other parameters fixed under L-ECM conditions, and conducted simulations for 10 cells over 600 minutes (**Fig. 4A**). Results for mechanical polarity, turning angle and speed of migration, number, maximum length, and lifetime of protrusions at *k*_*pr*_ = 0.2 are shown in **Fig. 4B-C** and ***SI Appendix*, Table S3**. As a result of the statistical tests (Wilcoxon rank-sum test), mechanical polarity decreased (p < 0.01). The migration speed decreased. Some cells in the simulation stopped moving, and the frequency of zero-speed events increased. However, no significant difference was observed in the turning angle. In contrast, the number of protrusions increased beyond the levels observed under fibronectin conditions, and the length decreased. Protrusion lifetimes also slightly decreased. We further varied *k*_*pr*_ from 0.1 to 0.2 in increments of 0.02, conducting simulations of 10 cells over 600 minutes for each condition, with data recorded every 2 minutes. The trends were largely similar. However, a significant increase in turning angle was observed (***SI Appendix*, Fig. S3**).

**Fig. 4.**
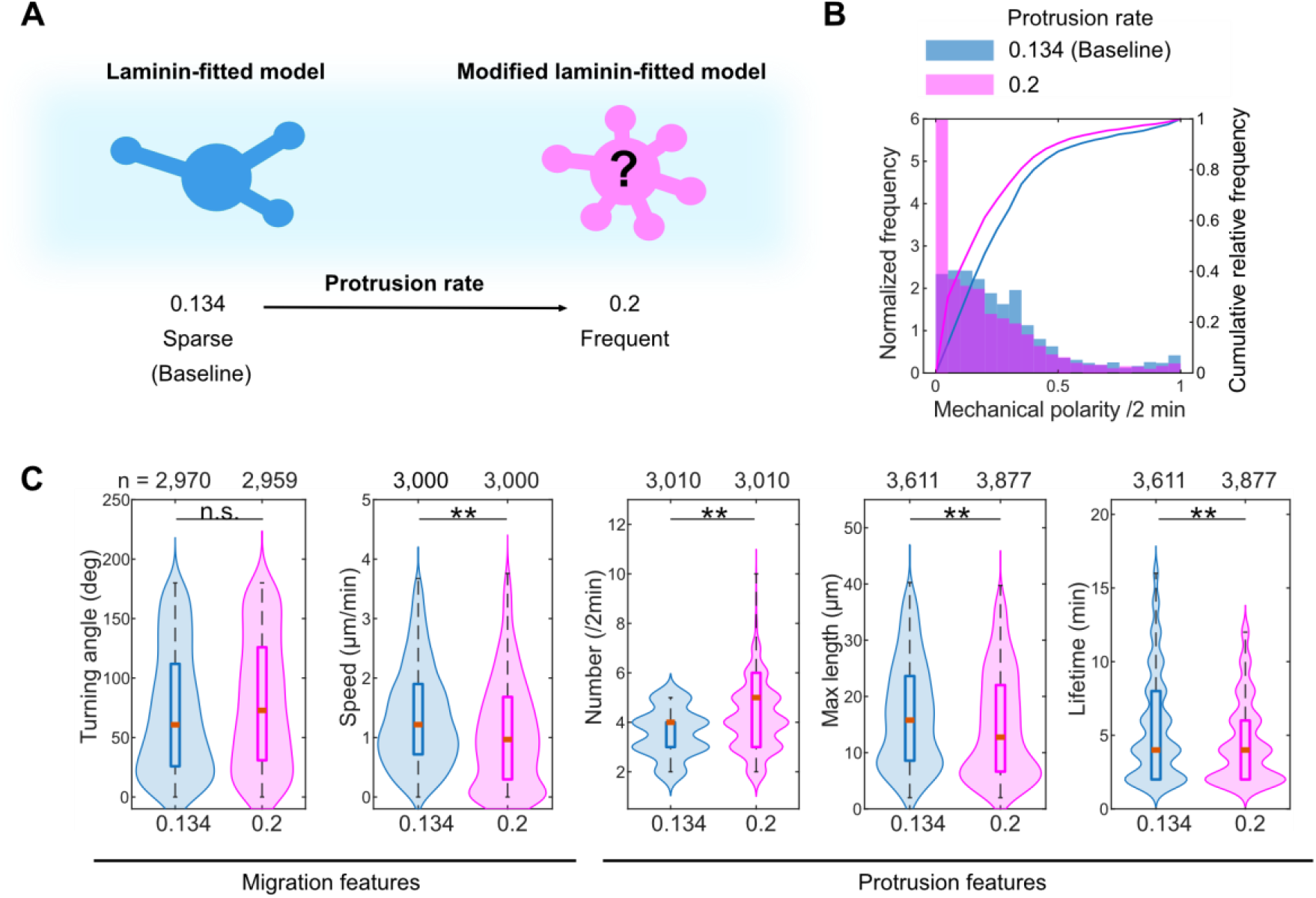
Effect of increased protrusion rate on cell motility. (A) Comparison of motility under two different values of the protrusion rate parameter *k*_*pr*_: baseline (*k*_*pr*_ = 0.134) and elevated (*k*_*pr*_ = 0.2). Parameters other than *k*_*pr*_ were set to the values under L-ECM conditions. (B-C) Comparison of motility features at *k*_*pr*_ = 0.134 (blue) and *k*_*pr*_ = 0.2 (magenta). Statistical differences were assessed using the Wilcoxon rank-sum test (**p < 0.01; n.s., not significant). The sample size *n* is indicated above each plot. The red line in the box represents the median. Values more than 1.5 times the interquartile range away from the top and bottom of the box were considered outliers. Outliers were omitted from the box plots for visibility, but all data were included in the statistical tests and analyses.

Since protrusions serve as the primary force generators for cell migration (27), the observed decrease in motility can be attributed to changes in protrusion features, specifically the increase in number and reduction in length. Collectively, these results suggest that both the protrusions and their mechanical balance play critical roles in regulating the efficiency of glioblastoma cell migration.

## Discussion

### Adhesion-tuned protrusions confer directional stability via mechanical polarity in cell migration

This study demonstrates that the motility of glioblastoma cells (GDC40) is strongly influenced by the type of ECM coating. Under laminin (L-ECM) conditions, cells exhibited fewer but longer protrusions, along with higher directional persistence and speed, whereas fibronectin (F-ECM) conditions led to more frequent, shorter protrusions and inefficient, random migration. Reconstruction using a mathematical model suggested that these differences arise from altered adhesion properties at protrusion tips, specifically changes in the bound fraction. These results indicate that ECM modulates motility through the physical parameters of adhesion, providing a way to reveal how a common internal mechanical system adapts to different boundary conditions.

Our model reproduced the observed migration behavior without invoking intracellular signaling, relying solely on the mechanical parameters of adhesion and detachment from the ECM. Traditionally, ECM-dependent differences in migration have been explained primarily by intracellular signaling (8, 28). Here, we showed that the migration efficiency we observed can be altered solely by differences in physical adhesion properties. Consistent with this view, neuronal axon outgrowth direction depends on differences in ECM adhesion (29), supporting our conclusion that physical adhesion properties can regulate cell motility.

Notably, the direction of cell body translocation is determined by the vector sum of tensile forces exerted by multiple protrusions. This phenomenon, in which directional migration emerges through mechanical coordination among protrusions, forms the basis for what we define as mechanical polarity. Rather than replacing morphological polarity, mechanical polarity complements it by providing directional stability even when morphological asymmetry is weak or transient. This framework offers a mechanical basis for understanding how highly plastic cells such as glioblastoma maintain persistent migration.

### Possible causes of ECM-dependent adhesion changes and how they contribute to protrusion remodeling and migration

The observed differences in adhesion dynamics between laminin and fibronectin likely originate from distinct molecular interactions at the cell–ECM interface. GDC40 cells express the L1 cell adhesion molecule (L1CAM), which has been reported to exhibit catch bond–like binding to laminin (30), and we confirmed that elevated cell surface expression of L1CAM enhances migratory capacity (Katsuma et al., submitted). In contrast, we are not aware of evidence for direct L1CAM–fibronectin binding. These are consistent with, and may underlie, the adhesion parameters estimated in our model.

The L1CAM–laminin interactions could stabilize adhesions under tensile load, enhancing force transmission and promoting the formation of longer, more persistent protrusions (27, 31). Longer protrusions tend to retain resources for elongation such as clutch molecules near the distal tip, as retrograde diffusion from the tip toward the cell body is limited, allowing certain protrusions to extend further. This resource-driven elongation mechanism has also been proposed as a driver of symmetry breaking in neuronal morphology (22). As a result, L-ECM conditions give rise to fewer but longer and more stable protrusions whose forces align toward the direction of migration.

By contrast, the absence of effective L1CAM–fibronectin interactions may result in weaker, more transient adhesions. This may cause the bound fraction to decrease more readily with increasing force (**Fig. 3A**), leading to shorter, unstable protrusions that fail to accumulate resources. Because each protrusion consumes fewer resources, more resources become available for new protrusions, resulting in a greater number of protrusions. These protrusions generate weak forces in multiple directions, which cancel out, reducing the net force bias that drives cell body displacement and thereby lowering migration efficiency.

Even if L1CAM does not bind to fibronectin, other molecules such as integrins do. Integrins are also known to exhibit catch bond with ECM ligands (18). They undergo force-induced conformational activation, enhancing binding to fibronectin (18, 25). However, integrins are known to bind to laminin (25) (but no catch bond to laminin has been observed). Thus, they may not contribute as much as L1CAM to the ECM-dependent differences. Future experimental quantification of these interactions will help bridge the mechanical and molecular levels of analysis, enabling more precise modeling of cell migration.

### Modulating protrusion emergence disrupts force coherence and suppresses migration

Increasing the protrusion emergence rate (*k*_*pr*_) alone, without altering adhesion dynamics, could reduce migration efficiency in our simulations. As the number of protrusions increased, force interference between them became more pronounced, diminishing the net driving force for migration. This relationship is illustrated in **Fig. 5**, which visually links protrusion patterns and force alignment to migratory behavior.

Molecular regulators such as the growth-associated protein 43 (GAP-43) (5) and tweety homolog 1 (Ttyh1) (32, 33) are known to promote tumor microtube protrusion. Testing whether the manipulation of these genes alters protrusion dynamics and migration efficiency in line with our model predictions could provide critical experimental validation.

**Fig. 5.**
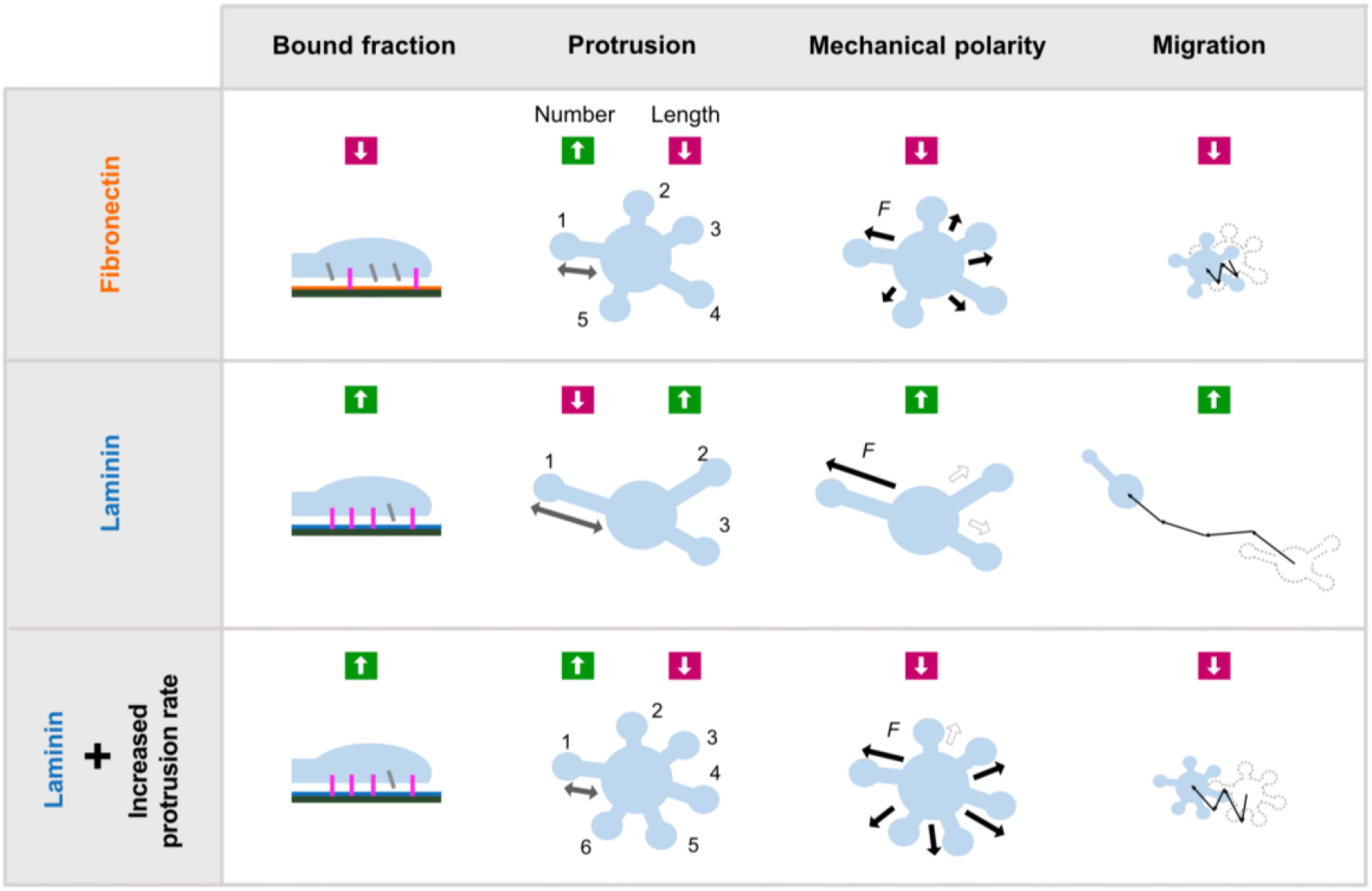
Conceptual model of how ECM and protrusion dynamics shape cell migration. Summary of migration-related features under three conditions: fibronectin (top), laminin (middle), and laminin with increased protrusion rate (bottom). Key characteristics (bound fraction, protrusion morphology, mechanical polarity, and overall migration) are represented schematically. In fibronectin conditions, weak adhesion leads to numerous short protrusions, resulting in force cancellation and impaired migration. In laminin, stronger adhesion with fewer, longer protrusions enables better force alignment and efficient migration. When the protrusion rate is artificially increased under laminin conditions, the system shifts toward reduced mechanical polarity and diminished migration.

### Future directions

The current model assumes homogeneous adhesion properties across all protrusions. In reality, each protrusion likely experiences different molecular compositions and mechanical environments. Extending the model to allow protrusion-specific bound fraction values would better reflect biological complexity. Variability in protrusion number, length, and lifetime (**Fig. 2C and D**) may also be captured by introducing stochastic gene expression fluctuations, such as in clutch molecules or cytoskeletal component levels.

Myosin II has been implicated in glioma invasion (34), and similar force-induced myosin II recruitment has also been reported in *Dictyostelium* (35). These observations suggest that in GDC40 cells, local tension might enhance myosin II accumulation and contractility, potentially creating a positive feedback loop that modulates the bound fraction. Such feedback could be incorporated into the model by making the parameters of the bound fraction dependent on myosin II concentration or activity.

Our model provides a physics-based quantitative framework for understanding glioblastoma cell motility. Its simplicity and extensibility make it suitable for large-scale parameter screening or personalized therapeutic design. Modulating parameters related to adhesion and protrusion dynamics may offer new opportunities to suppress tumor invasion.

## Materials and Methods

### Experimental procedure

In this study, we used time-lapse images of glioblastoma-derived cell line GDC40 in vitro. GDC40 is a cell line isolated and established from clinical samples of a glioblastoma patient (16). The culture of patient-derived cells for image acquisition was conducted at Osaka National Hospital in accordance with the principles of the Declaration of Helsinki, following approval from the Institutional Review Board of Osaka

National Hospital (approval number: 713). Written informed consent was obtained from the patient.

An IncuCyte ImageLock 96-well plate (Sartorius, Goettingen, Germany) was coated with laminin (Sigma; L2020 mouse, 1 mg/mL solution) at 2 μg/cm^2^ or fibronectin (Sigma; F1141 bovine plasma, 0.1% solution) at 5 μg/cm^2^ and incubated at 37 °C for 2 h, washed three times with PBS, and then used for analysis. The dissociated cells were seeded on a laminin- or fibronectin-coated IncuCyte ImageLock 96-well plate at a density of 3×10^2^ cells/well and cultured in the IncuCyte ZOOM live imaging system (Sartorius), which contained a 20× objective lens, at 37 °C in a 5% CO2 incubator. Time-lapse images were obtained every 2 min for 48 h.

For both laminin and fibronectin conditions, we analyzed five cells each. The spatial conditions of the ECM were uniform, and the conditions of ECM concentration, stiffness, and cell density were the same. The migration of each cell was recorded using hundreds of time-lapse images. The number of images per cell is listed in ***SI Appendix*, Table S4**. Videos of the cells on fibronectin and laminin are shown in **Movie S1 and S2**.

### Quantification of glioblastoma cell motility

We obtained coordinates of the cell body center and protrusion tips from time-lapse images of GDC40 on fibronectin (F-GDC40) and GDC40 on laminin (L-GDC40) (**Fig. 1A-C**). The cell body coordinates were defined as the center of the inscribed circle of the cell body. Protrusions were defined as local outward extensions of the cell contour when the cell body was approximated as a circle. Vertex-like points at angular edges of the cell perimeter were also classified as protrusions (see **Fig. 1A and C**). The protrusion tips were visually determined. Using these coordinates, we calculated five features: migration features (**Fig. 1D**), which include migration turning angle and speed, and protrusion features (**Fig. 1E**), which include the number of protrusions at each time point, the maximum length and lifetime of each protrusion. Turning angle was quantified using 4-minute intervals, which reduced noise caused by excessively short measurement intervals. Image processing and quantification were performed using MATLAB R2024a (MathWorks, Inc.) (***SI Appendix*, SI Text, Fig. S4 and S5**).

### Quantitative modeling of glioblastoma cell migration

#### Model overview

We constructed a migration model of glioblastoma cells (**Fig. 2A**) based on a previously developed mathematical model of neuronal polarization (22). In that model, neurons acquire protrusive forces at the tips via clutch molecules, which transmit the force of actin retrograde flow to the ECM. The protrusion length is determined by the mechanical balance between this protrusive force and tension. This mechanism, in which protrusion tips generate traction forces, is also relevant to glioblastoma cell motility (36).

In contrast to the neuronal model, however, the cell body of glioblastoma cells is not fixed but migrates as it is pulled by protrusions. Furthermore, in the neuronal model, the transmission of force via actin flow is regulated by the concentration of clutch molecules, while the force transmitted to the ECM is fixed and does not account for adhesion detachment. In this study, we extended the model by allowing the cell body to migrate in response to protrusive forces and by introducing an ECM-dependent force transmission function that permits detachment (***SI Appendix*, SI Text**). The full mathematical formulation of the model is presented below.

### Dynamics of cell body and protrusion tip

The glioblastoma cell body migrates by traction of protrusions (***SI Appendix*, Fig. S6A**). The protrusions are elongated when their tips bind to the ECM and exert stresses on the cell body (***SI Appendix*, Fig. S6B**). Assuming that inertial effects can be ignored because the environment in which the cell migrates is highly viscous, the position vector of the cell body center at time *t* (**x**_*cb*_(*t*)) is proportional to the sum of the forces:

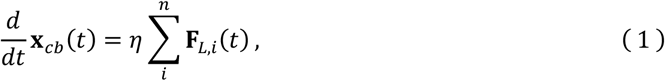

where **F**_*L*,*i*_(*t*) is the vector of tensile force of the *i*-th protrusion, parallel to the relative vector **L**(*t*) from the cell body center to the protrusion tip; *η* is the motility coefficient. As the cell body migrates according to this equation, the relative direction of each protrusion changes accordingly (***SI Appendix*, Fig. S7**), and **F**_*L*,*i*_(*t*) also changes.

From the definition of the vector above, the position vector of the protrusion tip at time *t* is the sum of **x**_*cb*_(*t*) and the relative vectors **L**(*t*) from **x**_*cb*_(*t*) to the protrusion tip:

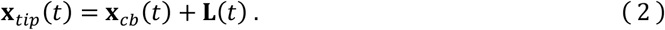

Its velocity has a direction parallel to the protrusion. It can be described as

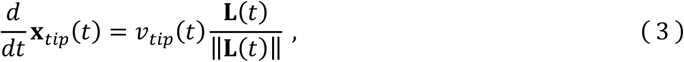

(***SI Appendix*, Fig. S7**).

### Model of elongation velocity at protrusion tip

Protrusions elongate through tubulin polymerization and retract through the disassembly of microtubules. The energy barrier for tubulin polymerization increases due to the load energy imposed by the membrane tension, and the membrane movement velocity is described by a force–velocity relationship (37, 38). Therefore, the membrane velocity at the protrusion tip is

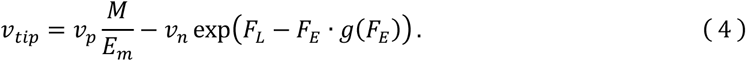

Here, *v*_*p*_ and *v*_*n*_ are polymerization and depolymerization constants, and *E*_*m*_ is the total tubulin amount (constant). *F*_*L*_ and *F*_*E*_ are variables related to the tension and protrusive force, respectively, and the function *g* represents the binding ratio of the adhesive molecules. For clarity, the protrusion index *i* and time *t* are omitted. The first term represents the polymerization rate dependent on the ratio of the time-varying tubulin monomer concentration *M* to *E*_*m*_, and the second term represents the disassembly rate based on the Arrhenius equation, which is force-energy dependent.

*F*_*L*_ represents the effect of nonlinear stress on microtubule depolymerization promotion due to protrusion elongation (dimensionless). Based on the previous study (22), this effect can be expressed as a monotonically increasing function of protrusion length *L*:

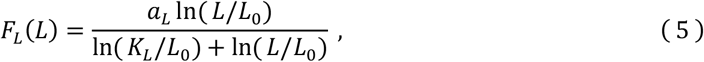

where *a*_*L*_ is the maximum value, *K*_*L*_ is the half-value constant, and *L*_0_ is the base length. This is a form of the first-order Hill equation applied to a logarithmic function, exhibiting linearity according to Hooke’s law when the protrusion is short. However, the stress increase amount decreases sharply as the protrusion lengthens to a certain extent (27).

*F*_*E*_ represents the effect of protrusive force on suppressing microtubule disassembly (dimensionless). It is a Hill-type increasing function of the clutch molecule concentration *C* (22):

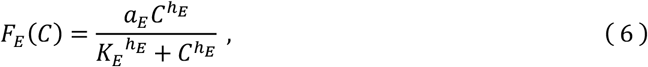

where *a*_*E*_ is the maximum value, *K*_*E*_ is the half-value constant, and *h*_*E*_ is the Hill coefficient.

*g*(*F*_*E*_) represents the bound fraction of adhesion molecules. We introduced this function into the glioblastoma cell migration model. It expresses the binding ratio of a population of adhesion molecules rather than a single adhesion molecule. The protrusive force by clutch molecules is generated when clutch molecules bind to adhesion molecules such as L1-CAM or integrin on the cell membrane. Adhesion molecules on the cell membrane repeatedly form catch bonds and slip bonds with ligands on the ECM (19, 39). In this study, we expressed the force transmitted via the clutch as the product of *g*(*F*_*E*_) and *F*_*E*_, representing the effective protrusive force effect.

We derived the bound fraction as a function of the clutch-induced protrusive force *F*_*E*_ based on previous

studies (19, 40) (***SI Appendix*, SI Text**):

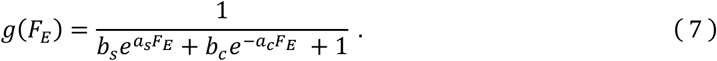

Here, *a*_*s*_ and *b*_*s*_ are parameters related to slip bonds, and *a*_*c*_, and *b*_*c*_ are parameters related to catch bonds. *g*(*F*_*E*_) takes values between 0 and 1 (**Fig. 2A-vi**). In the denominator, 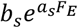 represents the reduction effect due to slip bonds, while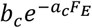 represents the increase effect due to catch bonds. The parameters were assumed to depend on the ECM.

### Protrusion emergence and retraction, and dynamics of cytoskeletal component amounts

Based on previous studies (41, 42), the protrusion emergence was probabilistically modeled. Under the condition that the amount of free tubulin *M* is greater than or equal to the amount required to form the initial protrusion length *L*_0_, the protrusion emergence probability per time step Δ*t* was modeled with a probability of

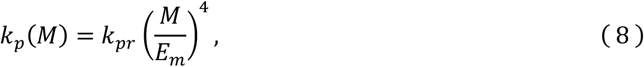

where *k*_*pr*_ is a coefficient representing the basic protrusion rate, and *E*_*m*_ denotes the total tubulin amount. Therefore, when the free tubulin ratio is small, the emergence probability decreases quadratically (41, 42). The direction of protrusion emergence is randomly determined around the cell body, and the emerged protrusions begin to elongate from the initial length *L*_0_. Simultaneously, *M* decreases by an amount equivalent to *L*_0_. Additionally, the condition for protrusions to retract is set as when the protrusion length becomes equal to or less than the initial length *L*_0_.

Free tubulin *M* increases or decreases according to the length *L*_*i*_ of each protrusion. When there are *n* protrusions, *M* is given by the following equation:

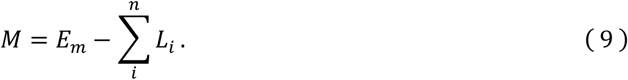

For clarity in the mathematical formulation, the amount of cytoskeletal components was rescaled to have the same unit as the protrusion length.

### Clutch molecule transport to protrusion tip and diffusion

Based on previous studies (22), the dynamics of clutch molecule concentration *C* at the tip of the protrusion were described as follows:

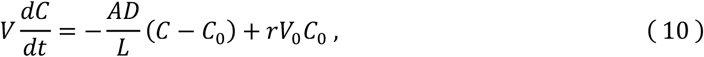

where constants *A, D, V*, and *V*_0_ represent the cross-sectional area of the protrusion, the diffusion coefficient, the volume of the protrusion tip and the cell body, respectively, and the remaining symbols are variables or functions. For clarity, the time *t* and protrusion index *i* are omitted. *C*_0_ is the clutch molecule concentration in the cell body, which changes while preserving the total amount *E*_*C*_ = *V*_0_ *C*_0_ + ∑_*i*_ *V*_*i*_*C*_*i*_.

The first term is a length-dependent diffusion term from the protrusion tip to the cell body based on the one-dimensional Fick’s law (43). The second term represents active transport from the cell body to the protrusion tip, which is determined by the transport rate *r* and *C*_0_. Lists of parameters, variables, and expressions used in this model are shown in ***SI Appendix*, Table S5-S7**.

### Parameter estimation and numerical simulation

This model contains about 6 more parameters than the neural polarization model due to changes such as introducing a bound fraction in the neural polarization model, resulting in a total of about 19 parameters. The estimation of these parameters was aimed at approximating the behavior of GDC40; both GDC40 and model glioblastoma cells have randomness in their motion and cannot be exactly matched to a specific individual motion. Therefore, the parameters were optimized so that the distributions of the quantitative features for the migration and protrusions of GDC40 (**Fig. 2C and D**) were similar to the corresponding distributions in the model. Note that some parameters were determined based on experimental data (***SI Appendix*, Fig. S8**) or previous studies (***SI Appendix*, Table S5**).

For this purpose, JS divergence was used as an indicator to evaluate the similarity between distributions, calculated for each of the five features, and the sum of these was used as the objective function. Parameter estimation was performed by minimizing this objective function (***SI Appendix*, SI Text**). The initial values of the parameters were set as follows: *E*_*C*_ = 2.5, *η* = 1.5, *h*_*E*_ = 2, *K*_*E*_ = 0.5, *a*_*E*_ = 10, *K*_*L*_ = 2.5, *v*_*p*_ = 550, *v*_*n*_ = 80, *k*_*pr*_ = 0.2, *a*_*L*_ = 2, *a*_*s*_ = 1.5, *b*_*s*_ = 0.001, *a*_*c*_ = 0.05, *b*_*c*_ = 1, *r V*_0_/*V* = 5.

In the numerical simulation, the time step was set to *dt* = 0.01 (min), and data on the migration and protrusions of 10 cells over a period of 600 minutes were collected as one set to create a distribution. At the start of the simulation, each cell was assigned three protrusions, with their generation directions determined randomly. The data from 60 minutes after the start to 660 minutes were used for analysis so that the period until cells began moving was excluded from the analysis. All simulations and analyses were performed using MATLAB R2024a (MathWorks, Inc.).

## Author Contributions

Conceptualization, Y.S., Y.K.;

Methodology, H.T., Y.S.;

Investigation, D.K., A.K.;

Resources, Y.K.;

Software, H.T.;

Validation, Y.K.;

Writing—original draft preparation, H.T.;

Writing—review and editing, Y.S., N.I., Y.K.;

Supervision, Y.K.;

Funding Acquisition, Y.S., Y.K., and N.I.;

All authors have read and agreed to the published version of the manuscript.

## Acknowledgments

We thank Prof. Yoichiro Hosokawa (NAIST) for his insightful discussions and continuous support throughout this study. We also acknowledge the laboratory members for their valuable technical assistance and constructive feedback. This work was partially supported by the Japan Society for the Promotion of Science (JSPS) KAKENHI (Grant Numbers 20H04283 and 23H04707 to Y.S.), the Japan Science and Technology Agency (JST) SPRING (Grant Number JPMJSP2140 to H.T.), and by the Japan Agency for Medical Research and Development (AMED) (Grant Number JP17gm0810011 to Y.S., Y.K., and N.I.).

## Data, Materials, and Software Availability

The data and software that support the findings of this study are available from the corresponding author upon reasonable request.

