## Supplementary material for "Mechanical polarity links adhesion-tuned protrusions to directional stability in glioblastoma cell migration": SI Appendix

### Supplementary Information for
### Mechanical polarity links adhesion-tuned protrusions to directional stability in glioblastoma cell migration

Haruna Tagawa[1#], Daisuke Kanematsu[2#], Asako Katsuma[2], Naoyuki Inagaki[1], Yonehiro Kanemura[2*], Yuichi Sakumura[1,3*]

1. Graduate School of Science and Technology, Nara Institute of Science and Technology, Ikoma 630-0192, Japan
2. Department of Biomedical Research and Innovation, Institute for Clinical Research, NHO Osaka National Hospital, Osaka 540-0006, Japan
3. Data Science Center, Nara Institute of Science and Technology, Ikoma 630-0192, Japan

[#]H.T. and D.K. contributed equally to this work.

**This PDF file includes:**

Supporting text
Figures S1 to S9
Tables S1 to S7
Legends for movies S1 to S4
SI References

**Other supporting materials for this manuscript include the following:**

Movies S1 to S4

#### Supplementary Information Text

#### Image processing procedure

##### 1. Acquisition of cell body center coordinates

The image processing procedures performed before manual correction of cell body coordinates are shown in **Fig. S4**. First, each frame in the original time-lapse sequence was normalized, followed by intensity inversion and binarization. Connected components smaller than 100 pixels were removed as noise, and holes within the segmented regions were filled to allow identification of the cell body as a single connected component. In cases where multiple cells appeared in the same image, the target cell body was manually selected in the first frame. In subsequent frames, the region with the greatest overlap with the previously selected area was tracked as the same cell. Within the segmented region of the target cell, the largest inscribed circle (1) was defined as the core region of the cell body, and its center was used as the cell body center. However, due to occasional missegmentation or displacement of the cell body region, we manually verified and corrected the center coordinates after this series of processing steps.

##### 2. Branch-like protrusion removal

In addition to protrusions extending from the cell body, branch-like protrusions can emerge from other protrusions. For simplicity, such "branch" structures were manually excluded from the analysis (**Fig. S5A**).

##### 3. Protrusion identification

To record the lifetime and the maximum length of each protrusion during its existence, it was necessary to identify individual protrusions over time. For this purpose, protrusions that appeared to be the same across consecutive image frames were assigned the same identification number (**Fig. S5B**).

Initial identification was performed automatically. The procedure, illustrated in **Fig. S5C**, involved comparing protrusion coordinates between consecutive frames—referred to as the previous image and the current image—to determine whether a given protrusion in the current image corresponded to one in the previous frame or represented a new one. The previous and current images were aligned based on the cell body center. For each candidate pair (a current protrusion and a protrusion in the previous image), two criteria were evaluated: (criterion 1) the distance between the coordinates near the base of the protrusion, and (criterion 2) the difference in the total number of coordinates, which approximates the protrusion length. Among the candidate protrusions in the previous image that met both criteria, the one with the smallest base distance (criterion 1) was considered to be the same protrusion and was assigned the same identifier. If no candidate in the previous frame satisfied both conditions, the current protrusion was treated as a new one and assigned a new identifier. This process was applied to each protrusion in each image. After automatic processing, manual inspection and corrections were performed as needed.

#### Additions and modifications to the neuronal polarity formation model for constructing the glioblastoma cell migration model

We constructed a cell migration model of GDC40 by adding to and modifying several elements of the neuronal polarity formation model (2). The six major additions and modifications are as follows.

##### 1. Non-probabilistic clutch molecule transport

Unlike neurons, probabilistic transport of clutch molecules to the tips of protrusions has not been observed in glioblastoma cells. Therefore, in glioblastoma cells, we introduced a deterministic molecular transport proportional to the concentration in the cell body, $C_0$. The dynamics of clutch molecule concentration at the protrusion tips $C$ in the glioblastoma cell model are as follows:

$$V\frac{dC}{dt} = -\frac{AD}{L}(C - C_0) + rV_0C_0 \, ,$$

where constants $A$, $D$, $V$, and $V_0$ represent the cross-sectional area of the protrusion, the diffusion coefficient, the volume of the protrusion tip, and the volume of the cell body, respectively.

##### 2. Constant total amount of cytoskeletal components and clutch molecules

In the neuronal polarity formation model, the expression levels of each molecule were changed over time based on the experimental data, as the model targeted neuronal polarity formation over several tens of hours. In this study, we focused on glioblastoma movement over 3–15 hours (**Table S1**), so the total amount of cytoskeletal components and clutch molecules in the cells was kept constant as $E_m$ and $E_C$ over time.

##### 3. Introduction of the bound fraction function

In this model, to take into account the adhesive properties with respect to the extracellular matrix (ECM), the product of the protrusive force function $F_E(C)$ and the function $g(F_E)$ representing the bound fraction was used as the net protrusive force:

$$F_E(C)\, g(F_E) = \frac{a_E C^{h_E}}{{K_E}^{h_E} + C^{h_E}}\, g(F_E)\,.$$

Derivation of $g(F_E)$ is described later.

##### 4. Movement of cell bodies due to pulling forces exerted by each protrusion

In the neuronal polarity formation model, the position of cell bodies was fixed. However, in this model, cell bodies were assumed to move due to pulling forces exerted by each protrusion (**Fig. S6**).

##### 5. Model of protrusion formation and disappearance, and dynamics of cytoskeletal molecular components

In the neuronal polarity formation model, the number of protrusions does not change. We added factors for protrusion formation and disappearance to this model to vary the number of protrusions over time.

##### 6. Modification of the dL/dt expression

Polymerization of cytoskeletal molecules is proportional to the amount of free cytoskeletal molecules $M$ relative to the total cytoskeletal pool $E_m$. Unlike in the neuronal polarity formation model, the position of the cell body is not fixed in the glioblastoma cell migration model. Therefore, we expressed polymerization as the change in the protrusion tip position $x_{tip}$ in absolute coordinates, rather than as the protrusion length $L$:

$$\frac{dx_{tip}}{dt} = v_{tip} = v_p \frac{M}{E_m} - v_n \exp\bigl(F_L - F_E \cdot g(F_E)\bigr)\ ,$$

where $v_p$ and $v_n$ are polymerization and depolymerization constants. $F_L$ is a variable related to the tension of the protrusion.

#### Derivation of the bound fraction function *g*

Assume that the total number of adhesion molecules on the cell side is $N_1$, the number of adhesion sites on the extracellular matrix side is $N_2$, and $N_b$ of these are bound. When $N_2 \gg N_b$, according to Bell et al. (6),

$$\frac{dN_b}{dt} = k_f(N_1 - N_b)N_2 - k_b N_b \exp\left(\frac{\gamma_s F}{K_B T N_b}\right).$$

Here, $k_f$ and $k_b$ are reaction rate constants ($s^{-1}$), $K_B$ is the Boltzmann constant ($m^2$ kg $s^{-1}$ $K^{-1}$), $T$ is the absolute temperature (K), $\gamma_s$ is a parameter determining the strength of the bond (μm) (7), and $F$ is the force acting on the binding site (N).

When the effect of catch bond (8, 9) is taken into account, it becomes

$$\frac{dN_b}{dt} = k_f(N_1 - N_b)N_2 - k_b N_b \exp\left(\frac{\gamma_s F}{K_B T N_b}\right) - k_{b,2} N_b \exp\left(\frac{-\gamma_c F}{K_B T N_b}\right).$$

Note that $\gamma_s > 0$ and $\gamma_c > 0$.

In a steady state $\left(\frac{dN_b}{dt} = 0\right)$,

$$0 = k_f(N_1 - N_b)N_2 - k_b N_b \exp\left(\frac{\gamma_s F}{K_B T N_b}\right) - k_{b,2} N_b \exp\left(\frac{-\gamma_c F}{K_B T N_b}\right),$$

and after reformulating the equation,

$$N_b = \frac{N_1}{\frac{k_b}{k_f N_2}\, exp\left(\frac{\gamma_s F}{K_B T N_b}\right) + \frac{k_{b,2}}{k_f N_2}\, \exp\left(\frac{-\gamma_c F}{K_B T N_b}\right) + 1}.$$

Here, if the bound fraction is $g$, $N_b = N_1\, g$, so

$$N_1 g = \frac{N_1}{\frac{k_b}{k_f N_2}\, \exp\left(\frac{\gamma_s F}{K_B T N_1 g}\right) + \frac{k_{b,2}}{k_f N_2}\, \exp\left(\frac{-\gamma_c F}{K_B T N_1 g}\right) + 1},$$

and

$$g = \frac{1}{\frac{k_b}{k_f N_2}\, \exp\left(\frac{\gamma_s F}{K_B T N_1 g}\right) + \frac{k_{b,2}}{k_f N_2}\, \exp\left(\frac{-\gamma_c F}{K_B T N_1 g}\right) + 1}.$$

If $F$ is the traction force $f_C\, g$ of the model (2), $F$ can be expressed as

$$F = f_C\, g = \frac{K_B T}{\delta} F_E\, g\,.$$

Here, $\delta$ is the length (μm) per unit of the skeletal components. Therefore,

$$g = \frac{1}{\frac{k_b}{k_f N_2} \exp\left(\frac{\gamma_s}{N_1\, \delta}\, F_E\right) + \frac{k_{b,2}}{k_f N_2} \exp\left(\frac{-\gamma_c}{N_1\, \delta}\, F_E\right) + 1}$$

$$= \frac{1}{b_{\mathrm{s}} \exp(a_{\mathrm{s}}\, F_E) + b_{\mathrm{c}} \exp(-a_{\mathrm{c}}\, F_E) + 1},$$

where

$$a_{\mathrm{s}} = \frac{\gamma_s}{N_1 \delta}, \qquad a_{\mathrm{c}} = \frac{\gamma_c}{N_1 \delta}, \qquad b_{\mathrm{s}} = \frac{k_b}{k_f N_2}, \qquad b_{\mathrm{c}} = \frac{k_{b,2}}{k_f N_2}$$

are used. For simplicity, the number of adhesion molecules $N_1$ on the cell side is assumed to be constant.

#### Calculation method for JS divergence

JS divergence (Jensen-Shannon divergence) is a metric that represents the difference between two distributions $p(x)$ and $q(x)$. When $p(x)$ and $q(x)$ are discrete distributions, it can be calculated using the following formula:

$$JS(p||q) = \frac{1}{2} KL(p\, ||\, R) + \frac{1}{2} KL(q\, ||\, R)\,, \tag{1}$$

where

$$R = \frac{p+q}{2}\,, \tag{2}$$

$$KL(p||R) = \sum_x p(x) \log\left(\frac{p(x)}{R(x)}\right) dx\ . \tag{3}$$

In this study, $p(x)$ represents the distribution obtained from simulation results, and $q(x)$ represents the distribution obtained from quantitative results of images. The JS divergence was used as an indicator of how well the simulation results reproduce the actual data. Note that even for distributions with continuous values, such as migration speed and directness, they were treated as discrete values by dividing them into intervals of $dx$. The bin width $dx$ was determined for each feature by applying the Freedman–Diaconis rule (11) to the distribution obtained from the images. The procedure for calculating the JS divergence between the distributions of the quantitative results from the images and the simulation results is as follows.

First, we normalized the distribution quantified from the simulation results and images by setting the bin width to $dx$ and approximating the distribution area to 1. We set the values of the bar heights at $x$ in each histogram as $p(x)$ and $q(x)$. We calculated the JS divergence between $p(x)$ and $q(x)$ using the above formula, targeting the range from the smaller lower limit to the larger upper limit of the two distributions (10). When $p(x) = 0$ or $q(x) = 0$, we replaced those values with 0.00001 (since the argument of the logarithm must be positive).

#### Supporting figures

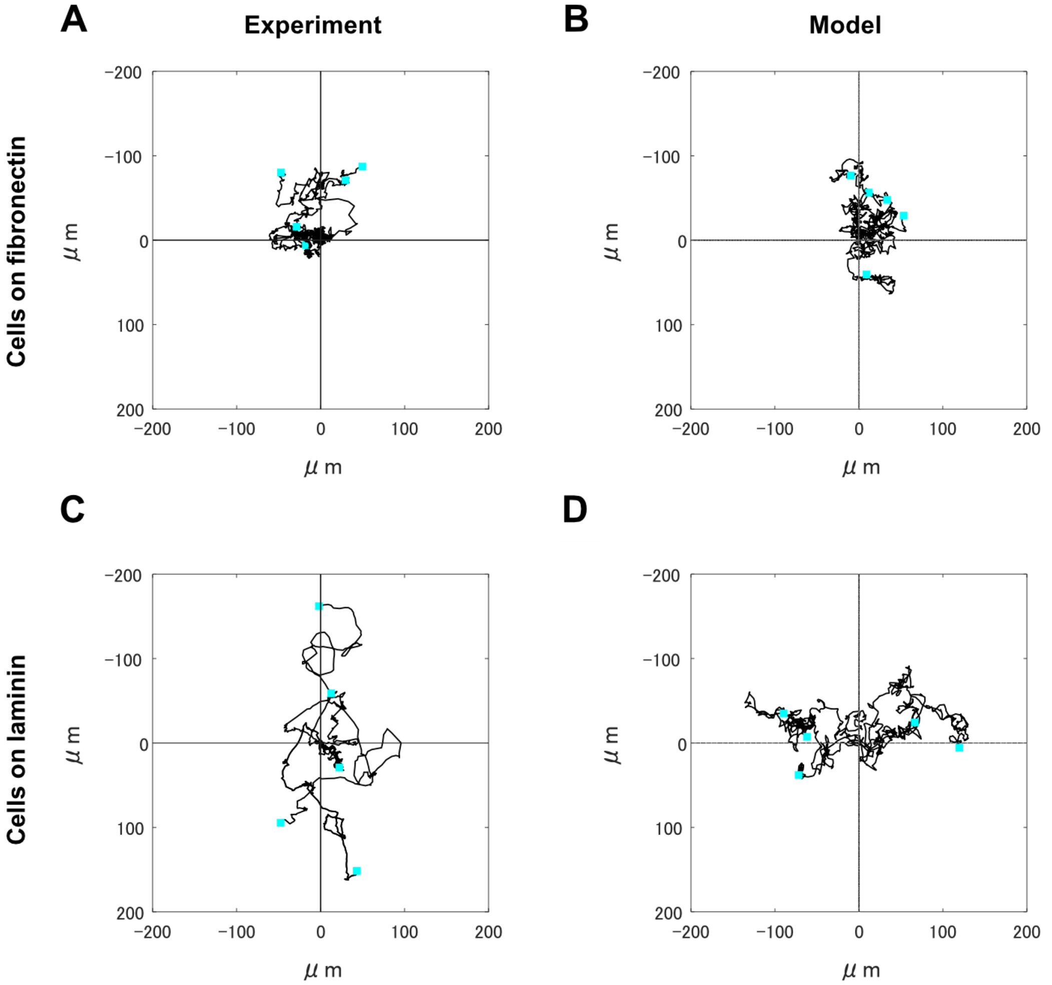


**Fig. S1. Cell migration trajectories.**

(A) Trajectories of cells migrating on fibronectin. (B) Simulated migration trajectories using parameters estimated from cell images on fibronectin (F-ECM). (C) Trajectories of cells migrating on laminin. (D) Simulated migration trajectories using parameters $a_s, b_s, a_c$ and $b_c$ estimated from cell images on laminin (L-ECM). Each panel shows data from five individual cells.

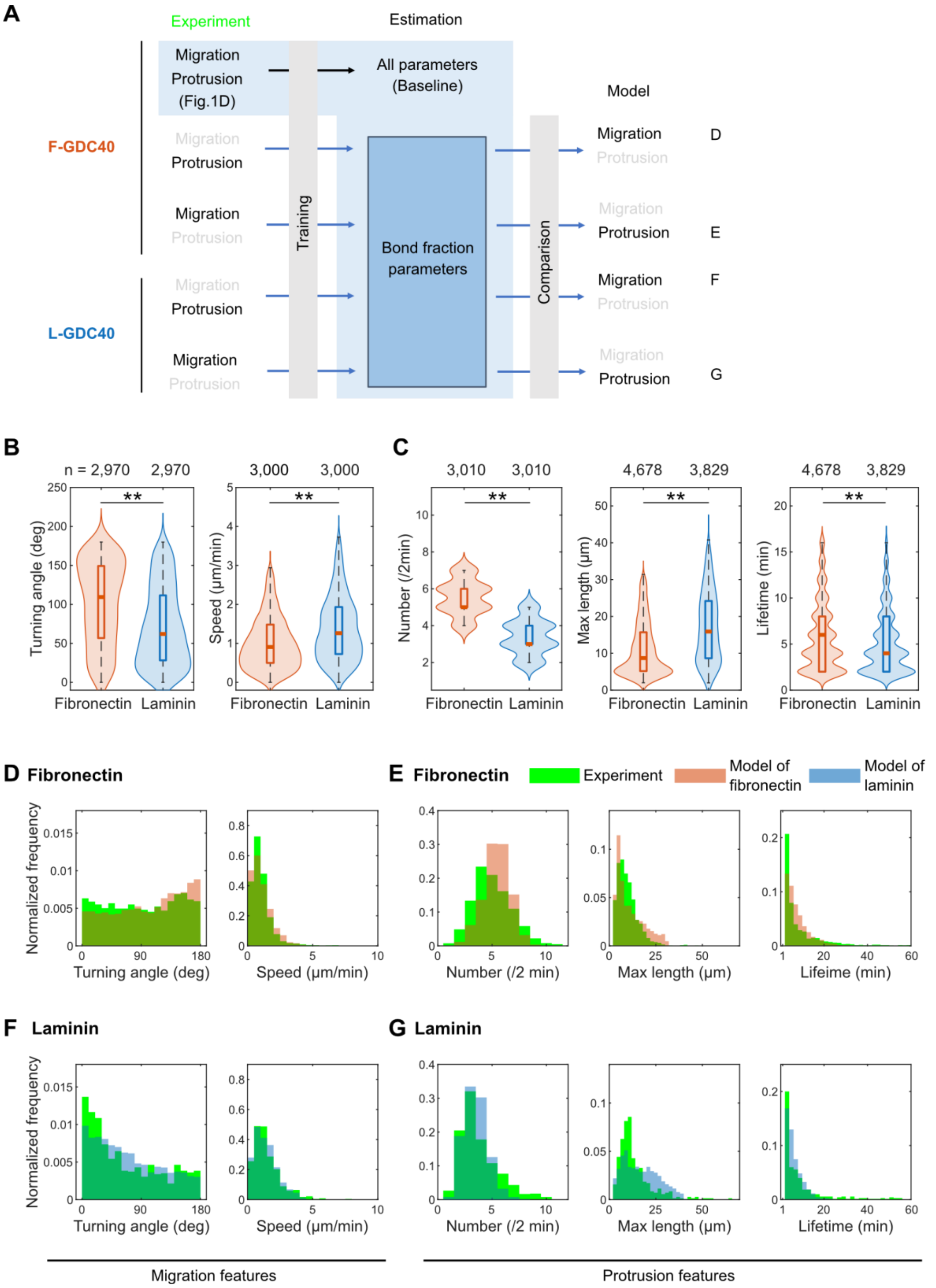


**Fig. S2. Reproduction of migration features from protrusion feature data, and vice versa.**

(A) Using the F-ECM condition as the baseline, bound fraction parameters $a_s$, $b_s$, $a_c$, and $b_c$ were estimated from the distribution of protrusion features (number, length, lifetime) for F-GDC40 or L-GDC40. The simulation results with the parameters are shown in (B), (D) and (F). Similarly, bound fraction parameters were estimated from the distributions of migration features (turning angle and speed) for F-GDC40 or L-GDC40. The simulation results with the parameters are shown in (C), (E) and (G). For comparison, the distributions obtained from actual cell images (green; same as in **Fig. 2C and D**) are also shown. Statistical differences were assessed using the Wilcoxon rank-sum test (**$p < 0.01$).

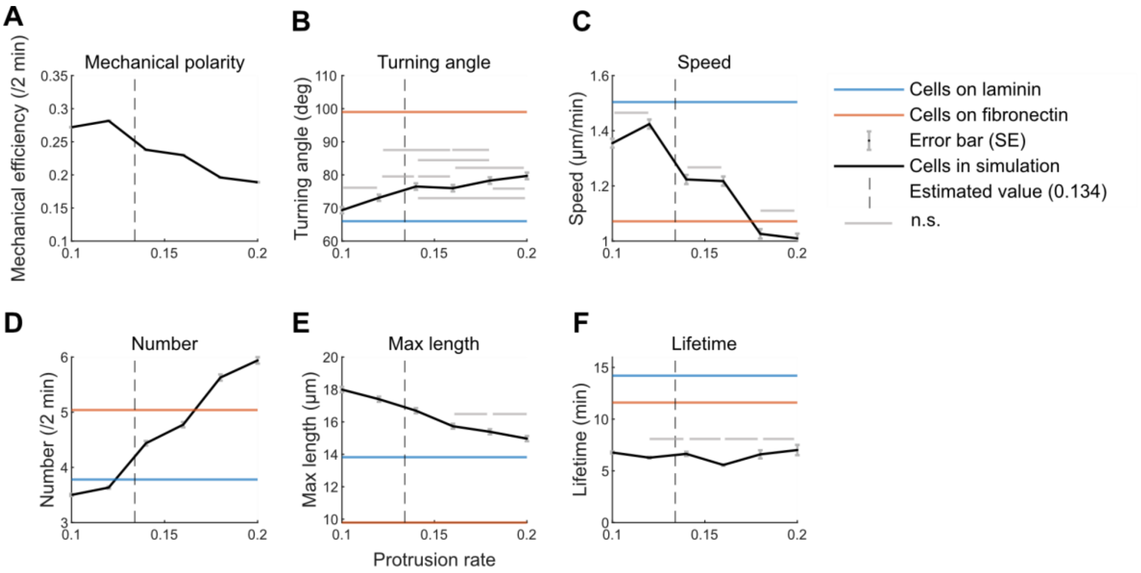


**Fig. S8. Sensitivity of migration features to protrusion rate coefficient $k_{pr}$**

Simulations were performed for $k_{pr}$ values from 0.1 to 0.2 (increment 0.02), with 10 cells simulated per condition for 600 minutes. Data were recorded every 2 minutes, and each panel shows mean ± standard error. (A–F) Mechanical polarity, turning angle, migration speed, protrusion number, maximum protrusion length, and lifetime, respectively. Orange and blue lines represent the experimental means of F-GDC40 and L-GDC40, respectively. The vertical dashed line indicates the fitted value of $k_{pr}$ for F-GDC40 ($k_{pr} = 0.134$). The gray horizontal lines indicate combinations that did not show significant differences in multiple-comparison tests (Tukey-Kramer method).

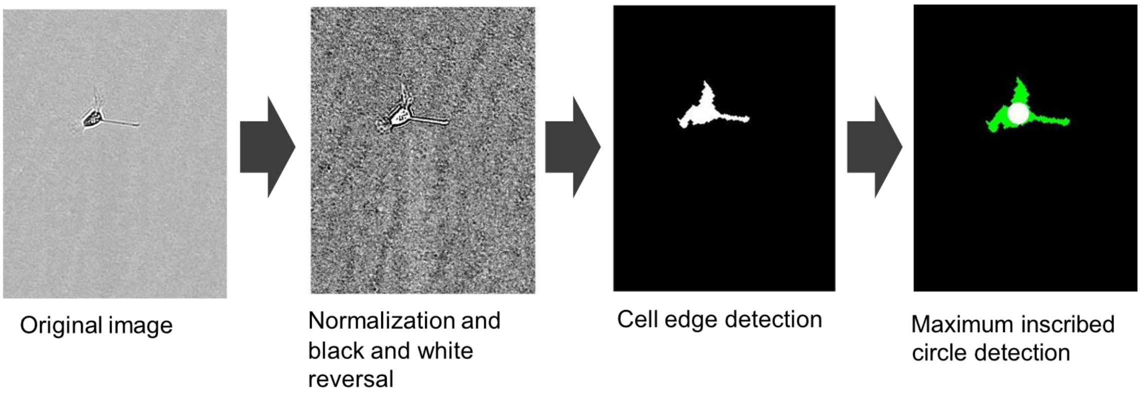


**Fig. S4. Method for automated extraction of cell body center coordinates.**
The original images were first normalized, followed by intensity inversion and binarization. Noise was removed, and holes were filled to enable identification of the target cell body region as a connected component. Since multiple cells may appear in a single image, the target cell was manually selected in the initial frame. In subsequent frames, the region with the greatest overlap with the previously selected region was tracked as the same cell. The center of the largest inscribed circle within the identified region was defined as the center of the cell body.

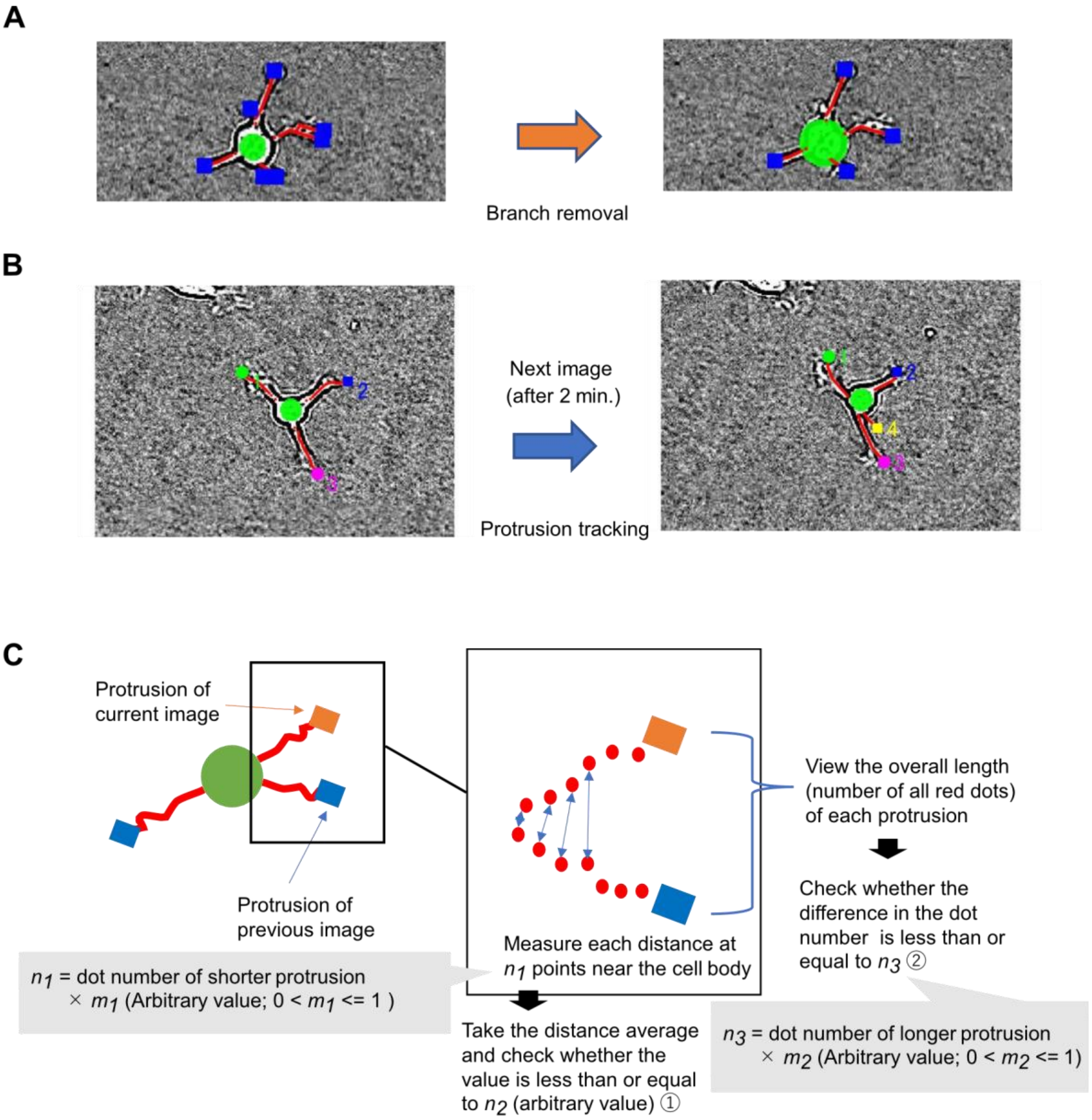


**Fig. S5. Processing of cellular protrusions.**

(A) Removal of branch-like structures. Protrusions that appeared to branch off from other protrusions, rather than directly extending from the cell body, were excluded from the analysis as branch-like structures. (B) Protrusion identification. The same identification number was assigned to protrusions that were judged to be the same across consecutive frames to identify individual protrusions across image frames. (C) Automatic method for protrusion identification. This panel illustrates how each protrusion in the current image (orange) was compared to protrusions in the previous image to determine whether it was the same or newly formed. The previous and current images were aligned based on the center of the cell body. For each pair consisting of a protrusion from the current image and one from the previous image, two conditions were evaluated: [1] whether the distances between the base coordinates near the cell body (specifically, the closest $n_1$ coordinates) were within a defined threshold $n_2$, [2] whether the difference in the total number of coordinates, which corresponds to protrusion length, was within a defined threshold $n_3$. Among the candidate protrusions in the previous image that met both conditions, the one with the smallest base distance was considered to be the same protrusion and was assigned the same identification number. If no candidate met both conditions, the protrusion was treated as a new one and assigned a new identifier. The parameter values used in this analysis were $n_2 = 25, m_1 = 0.8$, and $m_2 = 0.5$.

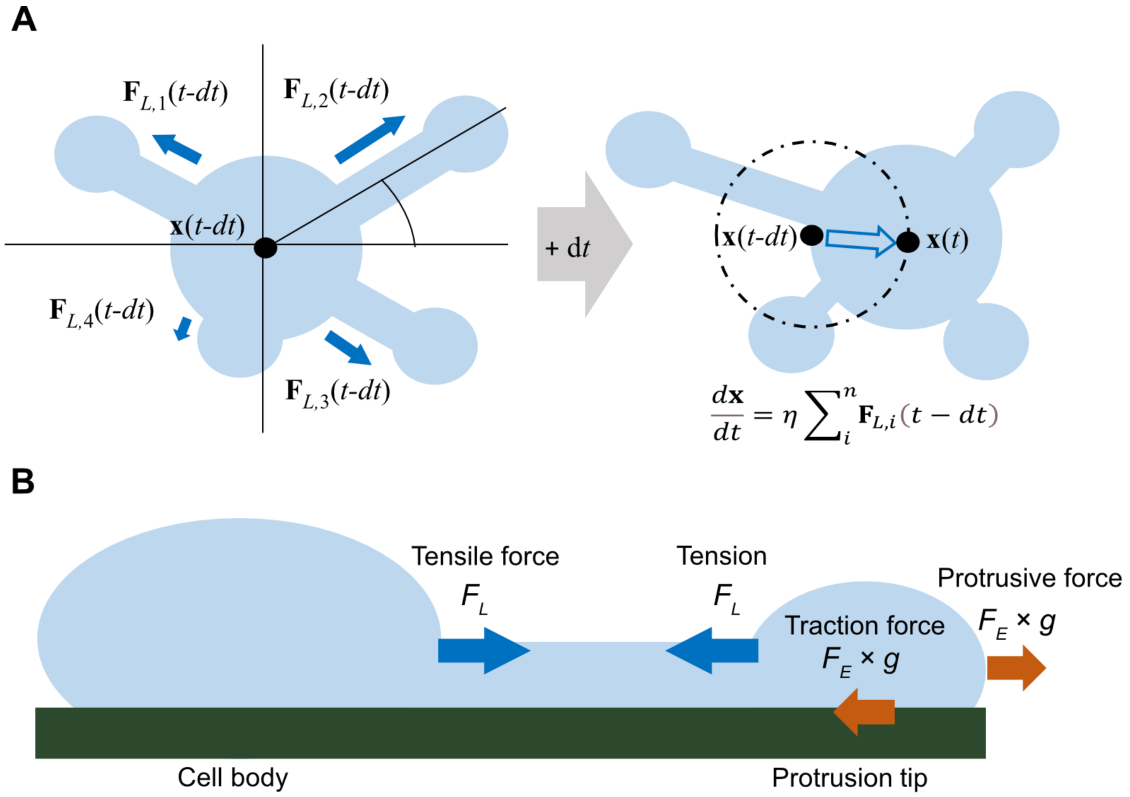


**Fig. S6. Displacement of the cell body position x due to protrusion-generated pulling forces.** (A) The cell body migrates in response to the tensile forces $\mathbf{F}_L$ generated by individual protrusions. The parameter $\eta$ is the motility coefficient associated with cell body movement. (B) While protrusion tip extension is driven by the protrusive force $F_E\ g$, which results from the traction force to substrate resistance via actin retrograde flow, the migration of the cell body is governed by the tensile force $F_L$ acting along the protrusion shaft. The tension applied during protrusion elongation promotes retraction.

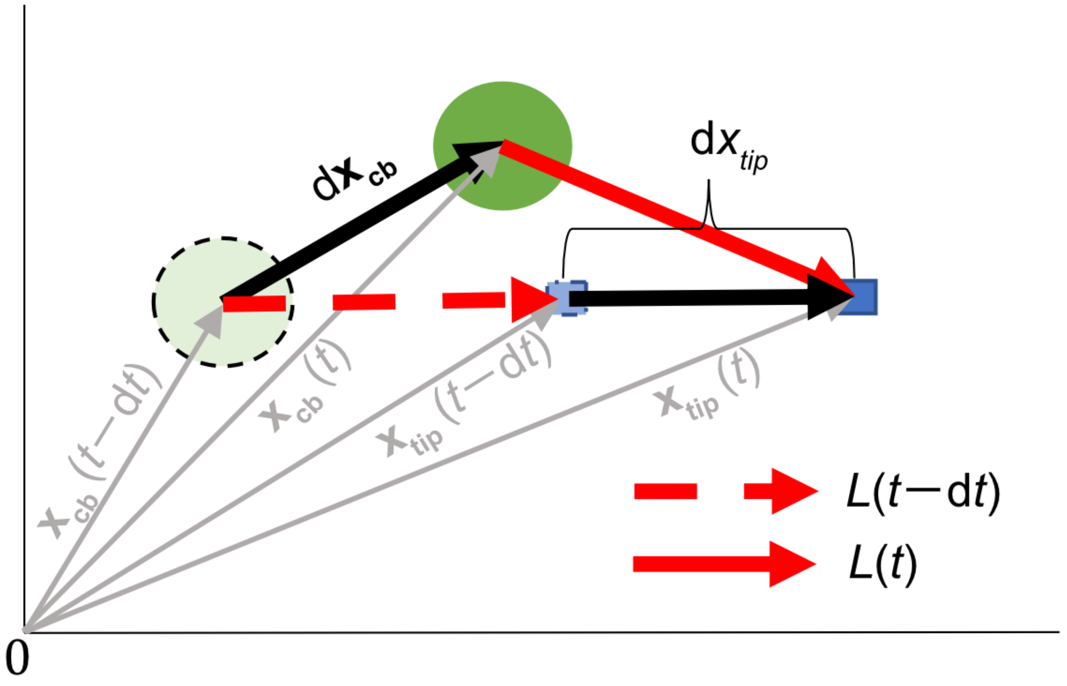


**Fig. S7. Cell body movement and protrusion elongation.**
At time point $t - dt$, $\mathbf{x}_{\mathrm{cb}}(t - dt)$ is the vector from the origin to the center of the cell body, and $\mathbf{x}_{\mathrm{tip}}(t - dt)$ is the vector from the origin to the tip of the protrusion. After a time interval $\mathrm{d}t$, if the protrusion elongates by $\mathrm{d}x_{tip}$, the vector from the origin to the tip becomes $\mathbf{x}_{\mathrm{tip}}(t)$. If the cell body does not move during this interval, the new protrusion length becomes the original length $L(t - 1)$ (indicated by the red dashed line) plus $\mathrm{d}x_{tip}$. However, if the cell body moves by $\mathrm{d}\mathbf{x}_{\mathrm{cb}}$, resulting in a new vector $\mathbf{x}_{\mathrm{cb}}(t)$, then the protrusion length is updated accordingly. The final length $L(t)$ is given by the magnitude of the updated vector from the new cell body center to the tip, as illustrated by the red solid line in the figure.

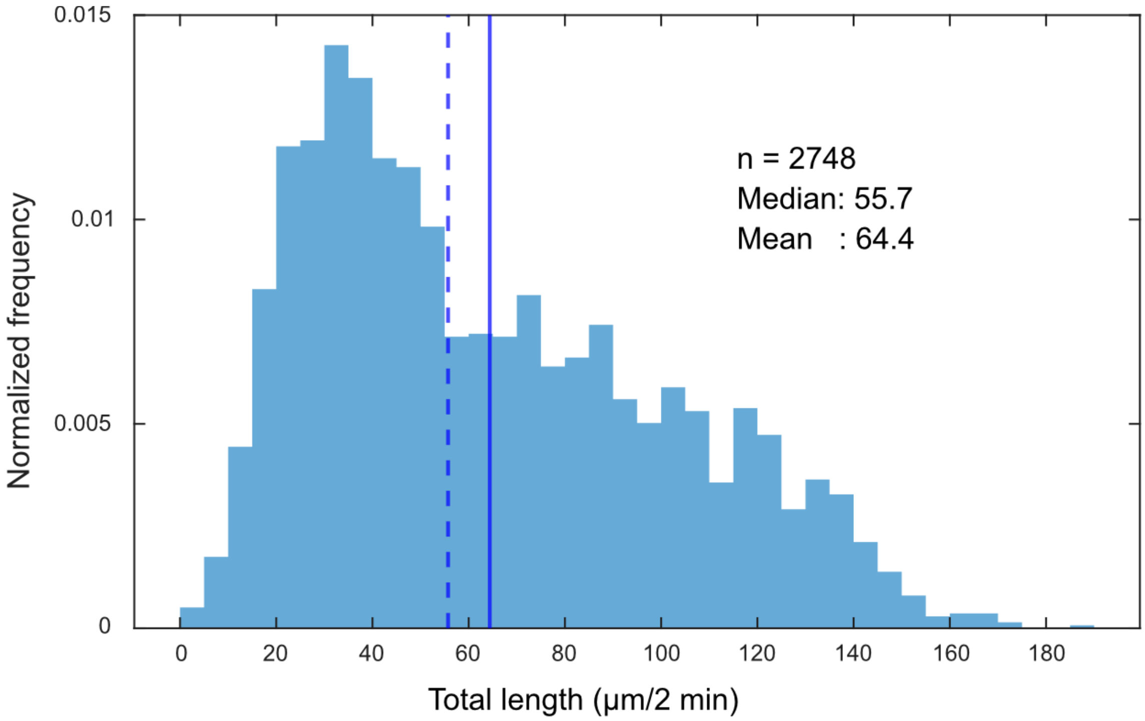


**Fig. S8. Total protrusion length per cell measured every 2 minutes in GDC40 cells.** Protrusion length was defined as the straight-line distance from the base to the tip of each protrusion. Results from 10 cells were compiled for both fibronectin and laminin ECM conditions. $n$ indicates the number of samples. The median (blue dashed line) and mean (blue solid line) are shown. In our cell migration model, the parameter $E_m$, representing the total length of cytoskeletal components within a cell, was set to the average of these total protrusion lengths (64.4 μm).

#### Supporting tables

**Table S1. Jensen–Shannon (JS) divergences comparing model feature distributions with experimental data under each ECM condition.**
Values are shown for the JS divergence between the F-ECM model distribution (orange in **Fig. 2C**) and the actual F-GDC40 distribution (green in **Fig. 2C**) or the actual L-GDC40 distribution (green in **Fig. 2D**), and the JS divergence between the L-ECM model distribution (blue in **Fig. 2D**) and the actual F-GDC40 distribution (green in **Fig. 2C**) or the actual L-GDC40 distribution (green in **Fig. 2D**).

| JS divergence of model vs experiment | Turning angle | Speed | Number | Max length | Lifetime |
|---|---|---|---|---|---|
| Fibronectin vs fibronectin | 0.009 | 0.018 | 0.031 | 0.046 | 0.039 |
| Fibronectin vs laminin | 0.012 | 0.026 | 0.134 | 0.049 | 0.053 |
| Laminin vs fibronectin | 0.020 | 0.042 | 0.098 | 0.120 | 0.046 |
| Laminin vs laminin | 0.006 | 0.010 | 0.046 | 0.055 | 0.061 |

**Table S2. JS divergences comparing model feature distributions with experimental data when adhesion parameters were estimated using only migration or protrusion features.**
Values are shown for the JS divergence between the F-ECM model distribution (orange in **Fig. S2D and E**) and the actual F-GDC40 distribution (green in **Fig. 2C**) or the actual L-GDC40 distribution (green in **Fig. 2D**), and the JS divergence between and the L-ECM model distribution (blue in **Fig. S2F and G**) and the actual F-GDC40 distribution (green in **Fig. 2C**) or the actual L-GDC40 distribution (green in **Fig. 2D**).

| JS divergence of model vs experiment | Turning angle | Speed | Number | Max length | Lifetime |
|---|---|---|---|---|---|
| Fibronectin vs Fibronectin | 0.010 | 0.016 | 0.049 | 0.034 | 0.044 |
| Fibronectin vs Laminin | 0.010 | 0.030 | 0.193 | 0.068 | 0.058 |
| Laminin vs Fibronectin | 0.020 | 0.047 | 0.132 | 0.123 | 0.052 |
| Laminin vs Laminin | 0.009 | 0.009 | 0.036 | 0.058 | 0.063 |

**Table S3. JS divergences comparing model feature distributions when the protrusion rate is increased.**
Values are shown for the JS divergence between the simulated distribution at $k_{pr} = 0.2$ (magenta in **Fig. 4D**) and the distribution for F-GDC40 (green in **Fig. 2C**) or the distribution for L-GDC40 (green in **Fig. 2D**).

| JS divergence of model ($k_{pr} = 0.2$) vs experiment | Turning angle | Speed | Number | Max length | Lifetime |
|---|---|---|---|---|---|
| vs fibronectin | 0.009 | 0.093 | 0.101 | 0.085 | 0.052 |
| vs laminin | 0.011 | 0.051 | 0.115 | 0.060 | 0.061 |

**Table S4. Cell labels and corresponding number of image frames.**
"Fib cell x" indicates a cell on fibronectin-coated substrate, and "Lam cell x" indicates a cell on laminin-coated substrate.

| Cell label | Number of images | Cell label | Number of images |
|---|---|---|---|
| fib cell 1 | 320 | lam cell 6 | 235 |
| fib cell 2 | 452 | lam cell 7 | 180 |
| fib cell 3 | 452 | lam cell 8 | 130 |
| fib cell 4 | 280 | lam cell 9 | 100 |
| fib cell 5 | 350 | lam cell 10 | 259 |

**Table S5. Summary of parameters used in the glioblastoma migration model.**
Parameters marked with an asterisk (*) were adopted from the original neuronal polarity formation model (2). The parameter $E_m$ was estimated from the average total protrusion length per time point, based on 2-minute interval time-lapse images (a total of 2,748 frames) from 10 glioblastoma cells (see **Fig. S8**). Parameters $r\,V_0/V\,, v_p, v_n, E_C, \eta, k_{pr}, a_E, h_E, K_E, a_L, K_L, a_s, b_s, a_c$ and $b_c$ were estimated from the distribution of quantitative features obtained from GDC40 cells migrating on fibronectin (F-ECM condition; see main text). The parameters $a_s, b_s, a_c$ and $b_c$ were also estimated from data on laminin-coated substrates (L-ECM condition; see main text).

| Parameter | Unit | Value | Meaning |
|---|---|---|---|
| $A\,D\,/\,V$ | μm/min | 8.26 * | Effective diffusion coefficient per unit length. $A$, $D$, and $V$ represent the cross-sectional area of the protrusion, the diffusion coefficient of the clutch molecule, and the volume of the protrusion tip, respectively. |
| $r\,V_0/V$ | /min | 4.88 | The ratio ( $r$ ) of clutch molecules transported per unit time from the cell body, and the coefficient for concentration conversion based on the volume ratio between the cell body and the protrusion tip ($V\,/V_0 = 1/10$). |
| $v_p$ | μm/min | 508 | Polymerization rate constant of cytoskeletal components |
| $v_n$ | μm/min | 52.1 | Depolymerization rate constant of cytoskeletal components |
| $E_m$ | μm | 64.4 | Total amount of cytoskeletal components in whole cell |
| $E_C$ | Relative conc. | 1.48 | Total amount of clutch molecules in whole cell |
| $\eta$ | μm/min | 1.15 | Cell motility coefficient |
| $k_{pr}$ | - | 0.134 | Coefficient at protrusion rate $k_p$ |
| $a_E$ | - | 6.54 | Maximum value of $F_E$ |
| $h_E$ | - | 2.04 | Hill coefficient |
| $K_E$ | Relative conc. | 0.595 | Half-value constant of $F_E$ |
| $a_L$ | - | 1.71 | Maximum value of $F_L$ |
| $K_L$ | μm | 2.07 | Half-value constant of $F_L$ |
| $L_0$ | μm | 2 | Natural protrusion length and reference length at the time of retract |
| $a_s$ | - | F-ECM: 2.00<br>L-ECM: 1.87 | Parameter of function $g$ |
| $b_s$ | - | F-ECM: $8.54\times10^{-4}$<br>L-ECM: $6.19\times10^{-4}$ | Parameter of function $g$ |
| $a_c$ | - | F-ECM: $3.70\times10^{-2}$<br>L-ECM: 0.040 | Parameter of function $g$ |
| $b_c$ | - | F-ECM: 1.21<br>L-ECM: 1.03 | Parameter of function $g$ |

**Table S6. Variables in the glioblastoma migration model**

| Variable | Unit | Meaning |
|---|---|---|
| $C_i$ | Relative conc. | Concentration of clutch molecules at the tip of protrusion $i$ |
| $C_0$ | Relative conc. | Concentration of clutch molecules in the cell body |
| $M$ | μm | Total amount of cytoskeletal components (total length) |
| $L_i$ | μm | Length of protrusion $i$ |
| $t$ | min | Time in simulation |

**Table S7. Expressions in the glioblastoma migration model**

| Formula | Meaning |
|---|---|

| | |
|---|---|
| $v_{tip} = v_p \frac{M}{E_m} - v_n \exp\big(F_L - F_E \cdot g(F_E)\big)$ | Speed of protrusion elongation at the protrusion tip |
| $F_E(C) = \frac{a_E C^{h_E}}{{K_E}^{h_E} + C^{h_E}}$ | Effect of protrusive force on the concentration of clutch molecules at the protrusion tip |
| $F_L(L) = \frac{a_L \ln(L/L_0)}{\ln(K_L/L_0) + \ln(L/L_0)}$ | Effect of stress on protrusion elongation depending on protrusion length |
| $g(F_E) = \frac{1}{b_s e^{a_s F_E} + b_c e^{-a_c F_E} + 1}$ | Bound fraction: the extent to which adhesion molecules at the protrusion tip are bound to the substrate |
| $\frac{dC}{dt} = -\frac{AD}{VL}(C - C_0) + r\frac{V_0}{V} C_0$ | Change in the concentration of clutch molecules at the protrusion tip |
| $\frac{d\mathbf{x}_{cb}}{dt} = \eta \sum_i^n \mathbf{F}_{L,i}(t - dt)$ | Change in the position of the cell body $\mathbf{x}_{cb}$: determined by the pulling force of each protrusion |
| $k_p(M) = k_{pr}\left(\frac{M}{E_m}\right)^4$ | Protrusion emergence rate: proportional to the fourth power of the amount of cytoskeletal components (12, 13) |
| $M = E_m - \sum_i^n L_i$ | Free cytoskeletal component amount (total length): defined as the total amount of components $E_m$ minus the length of each protrusion $L_i$ |
| $L(t) = \lvert\mathbf{x}_{tip}(t) - \mathbf{x}_{cb}(t)\rvert$ | Length of protrusion $i$ : defined as the distance from the cell body center to the protrusion tip |

#### Movie Legends

**Movie S1 (separate file). Cell migration on fibronectin**
GDC40 migration on a 2D plane on a fibronectin-coated matrix. The matrix concentration gradient was uniform.

**Movie S2 (separate file). Cell migration on laminin**
GDC40 migration on a 2D plane on a laminin-coated matrix. The matrix concentration gradient was uniform.

**Movie S3 (separate file). Cell migration in simulation (F-ECM condition)**
Model simulation video assuming cell migration on fibronectin. Parameters estimated from images of GDC40 cell migration on fibronectin were used (F-ECM condition).

**Movie S4 (separate file). Cell migration in simulation (L-ECM condition)**
Model simulation video assuming cell migration on laminin. Parameters estimated from images of GDC40 cell migration on laminin were used (L-ECM condition).
